# Real-time decoding of 5 finger movements from 2 EMG channels for mixed reality human-computer interaction

**DOI:** 10.1101/2021.09.28.462120

**Authors:** Eric J. McDermott, Thimm Zwiener, Ulf Ziemann, Christoph Zrenner

## Abstract

The search for optimized forms of human-computer interaction (HCI) has intensified alongside the growing potential for the combination of biosignals with virtual reality (VR) and augmented reality (AR) to enable the next generation of personal computing. At the core, this requires decoding the user’s biosignals into digital commands. Electromyography (EMG) is a biosensor of particular interest due to the ease of data collection, the relatively high signal-to-noise-ratio, its non-invasiveness, and the ability to interpret the signal as being generated by (intentional) muscle activity. Here, we investigate the potential of using data taken from a simple 2-channel EMG setup to differentiate 5 distinct movements. In particular, EMG was recorded from two bipolar sensors over small hand muscles (*extensor digitorum*, *flexor digitorum profundus*) while a subject performed 50 trials of dorsal extension and return for each of the five digits. The maximum and the mean values across the trial were determined for each channel and used as features. A k-nearest neighbors (kNN) classification was performed and overall 5-class classification accuracy reached 94% when using the full trial’s time window, while simulated real-time classification reached 90.4% accuracy when using the constructed kNN model (k=3) with a 280ms sliding window. Additionally, unsupervised learning was performed and a homogeneity of 85% was achieved. This study demonstrates that reliable decoding of different natural movements is possible with fewer than one channel per class, even without taking into account temporal features of the signal. The technical feasibility of this approach in a real-time setting was validated by sending real-time EMG data to a custom Unity3D VR application through a Lab Streaming Layer to control a user interface. Further use-cases of gamification and rehabilitation were also examined alongside the integration of eye-tracking and gesture recognition for a sensor fusion approach to HCI and user intent.

## Introduction

Electromyography (EMG) is the standard methodology used to measure muscle activity (Bigland-Ritchie 1981), and it has been used in controlling prosthetic devices (Castellini et al. 2009; Cipriani et al. 2008; Kiguchi et al. 2004; Kim et al. 2019), robotics (Fukuda et al. 2003; Gruver 1994) and rehabilitation (Graupe et al. 1982; Kawase et al. 2017). William Putnam was one of the first to describe a real-time EMG system (1993), and now, EMG and other wrist-based modalities are being explored as an input device for virtual, augmented, mixed, and extended reality applications for both research (Aung et al. 2011; Chowdhury et al. 2013; Jung et al. 2013; Kwon et al. 2020; Viriyasaksathian et al. 2011) and commercial purposes (Pezent et al. 2019; Young et al. 2019). The addition of biosignal-based agency within these mixed realities can help close the action-confirmation loop and offer critical feedback to improve performance and increase immersion (Sigrist et al. 2013). Finger movements already serve as a base control input in learned motor patterns such as mouse clicking, keyboard typing, and button pushing, yet it remains a technological challenge how such biologically based behaviors can be seamlessly integrated into the virtual world without manipulatable tools (i.e., mouse and keyboard) in order to increase human agency and immersion.

One approach at this “integration” requires a reliable translation of different movement-based actions into distinct classes of input based on the biological signals gathered from the actions, in a machine learning based procedure known as “decoding”. The first part consists of extracting a small number of features from the raw EMG signal, while the second part consists of determining the probability that the observed features were generated by a specific movement. In supervised learning, this requires use of a classification algorithm which is calibrated using a subset of the total data as training data. A number of previous studies have presented methodologies implementing this approach (Englehart et al. 2001; Hudgins et al. 1993; Zardoshti-Kermani et al. 1995), some achieving up to 93% on an online four-class problem (thumb, index, middle, hand closed) using 4 distinct EMG channels (one channel per digit) (Tsenov et al. 2006).

Here, we investigate the potential of achieving high classification accuracy of individual finger movements with minimal hardware cost and demands while using a simple feature extraction method. Specifically, we use only 2 distinct EMG channels and do not consider temporal features of the signal. Furthermore, we explore the potential to integrate this into a portable off-the-shelf real-time system by utilizing C++ based serial streaming from an Arduino Uno board (http://www.arduino.cc/), into a Python based model (http://www.python.org/), and then into a C# based Unity3D application (http://www.unity3d.com/) using a Lab Streaming Layer (Kothe et al. 2012). The use of EMG in real-time allows for versatile interaction with computer interfaces, and can be further optimized by combining it with eye-tracking as a second-level check of user intention (i.e., gaze fixation). This concept of *sensor fusion* can also be further extended by the inclusion of integrated gesture recognition given advances in computer vision.

## Experimental Methods

Data was collected through a serial stream coming from the Arduino Uno board via USB connection. A commercially available EMG sensor (“Myoware Muscle Sensor”) was used to extract the envelope of the EMG signal at 10-bit resolution, sampled at a default rate of 70 Hz. Sensors were placed at both the *extensor digitorum* (ED) and *flexor digitorum profundus* (FD). No filters were applied. Data for this single-subject proof-of-principle study was collected from one of the experimenters. First, the subject placed their arm and hand to rest comfortably on a table, with the wrist flat and the elbow padded. Then, the subject was instructed in a short calibration phase to perform a ‘maximum’ muscle contraction for each of the two EMG electrode sites (i.e., active wrist extension for ED, and active wrist flexion for FD), and a ‘minimum’ muscle contraction (i.e., be completely relaxed). These values then defined the subject’s range of output values, and the subsequently recorded data were normalized accordingly within these limits. No incoming data values exceeded the established minimum or maximum.

The main measurement consisted of 50 repetitions of the sequential active dorsal extension of each individual digit and subsequent return to the baseline rest position over a 2-second window, in an up-and-down motion, with 2-second pauses in between each movement. The beginning of each trial was cued by a countdown presented on a computer monitor.

To account for reaction time delays (on average between 150-400ms according to previous studies (Kutas et al. 1980; Ladd 1911; Murakami 2010)), and other edge effects, the first 200ms and last 200ms of each trial were removed, resulting in trials 1600ms long. Then, for each electrode, the max and mean value of the EMG envelope for each of the 50 trials was calculated, as well as for sliding windows of 70, 140, 280, 560, and 1020 milliseconds of each trial. The feature space was explored by a grid search of different variations of these features until the optimal feature set was found for both complete trial prediction and real-time prediction. Logistic, support vector machine (SVM), k-nearest neighbors (kNN), and DBSCAN (Hahsler et al. 2019) classifiers were compared and tested.

To validate the approach in a real-time setting, the resulting machine learning model was saved, and then a serial read was called from Python, where the incoming single time point data was transformed into a sliding average and sliding max. These newly created variables were then used as input features. Next, the resultant output class was directly sent to Unity3D using a Lab Streaming Layer. The virtual environment was designed and navigated with the HTC Vive (http://www.vive.com), furthermore the integrated eye-tracking from SRanipal (https://developer.vive.com/resources/vive-sense/sdk/vive-eye-tracking-sdk-sranipal/) was used to affirm gaze locking on an object, and if confirmed, EMG activity could then manipulate the object or otherwise interface with it accordingly. To further extend the amount of use cases for this paradigm, additional gesture recognition using the HTC Vive Hand Tracking SDK (https://developer.vive.com/resources/vive-sense/sdk/vive-hand-tracking-sdk/) was also applied in particular instances, such as to instantiate interactable user interfaces.

## Results

### Supervised Learning Approach

To visualize the relationship between EMG sensor activation and finger extension, a kernel density estimation with a Gaussian kernel was performed to obtain the probability density of the distribution of EMG sensor values received during the trials for each of five digits, for each EMG sensor, as shown in **Fig 1.**

**Fig 1:**
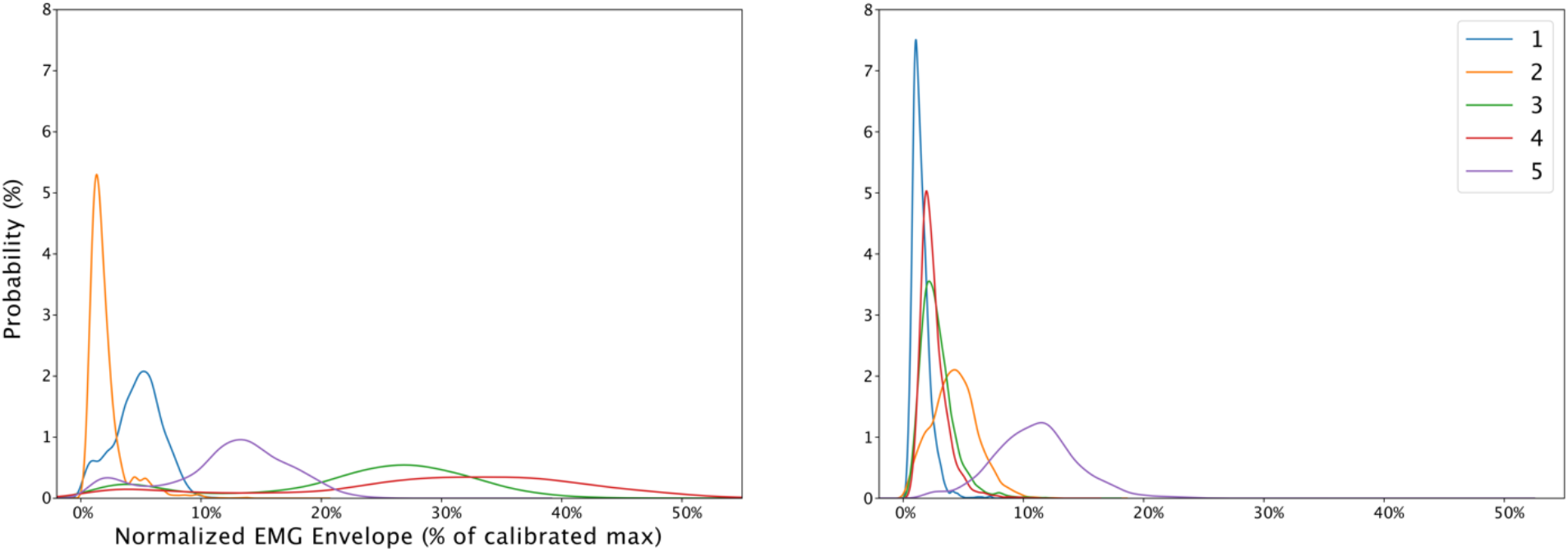
Kernel density estimation using Gaussian kernel for muscle activations in the ED (left panel) and FD (right panel) muscles. The horizontal axis corresponds to the EMG envelope (as a percentage of the maximum calibrated strength). Digits are numbered and color coded in sequential order starting with the thumb.

For the subsequent classification, the ‘mean’ and ‘max’ for each of the two EMG channels during each digit’s movement was extracted, obtaining four features, see **Fig 2a**. Substantial separation between classes can be seen in this four-dimensional feature-space by visual inspection. However, because determining the *max* or *mean* value for a whole trial requires the entire time window, this classification can only be performed *after* and not during the trial, thereby restricting its usage for real-time applications. In order to enable classification *during* finger extension, the same ‘mean’ and ‘max’ features were also extracted from a sliding window with a length of 280ms (20 data points), see **Fig 2b**, which is near the upper limit of latency still enabling a responsive user experience for basic applications (Henderson 2001).

**Fig 2:**
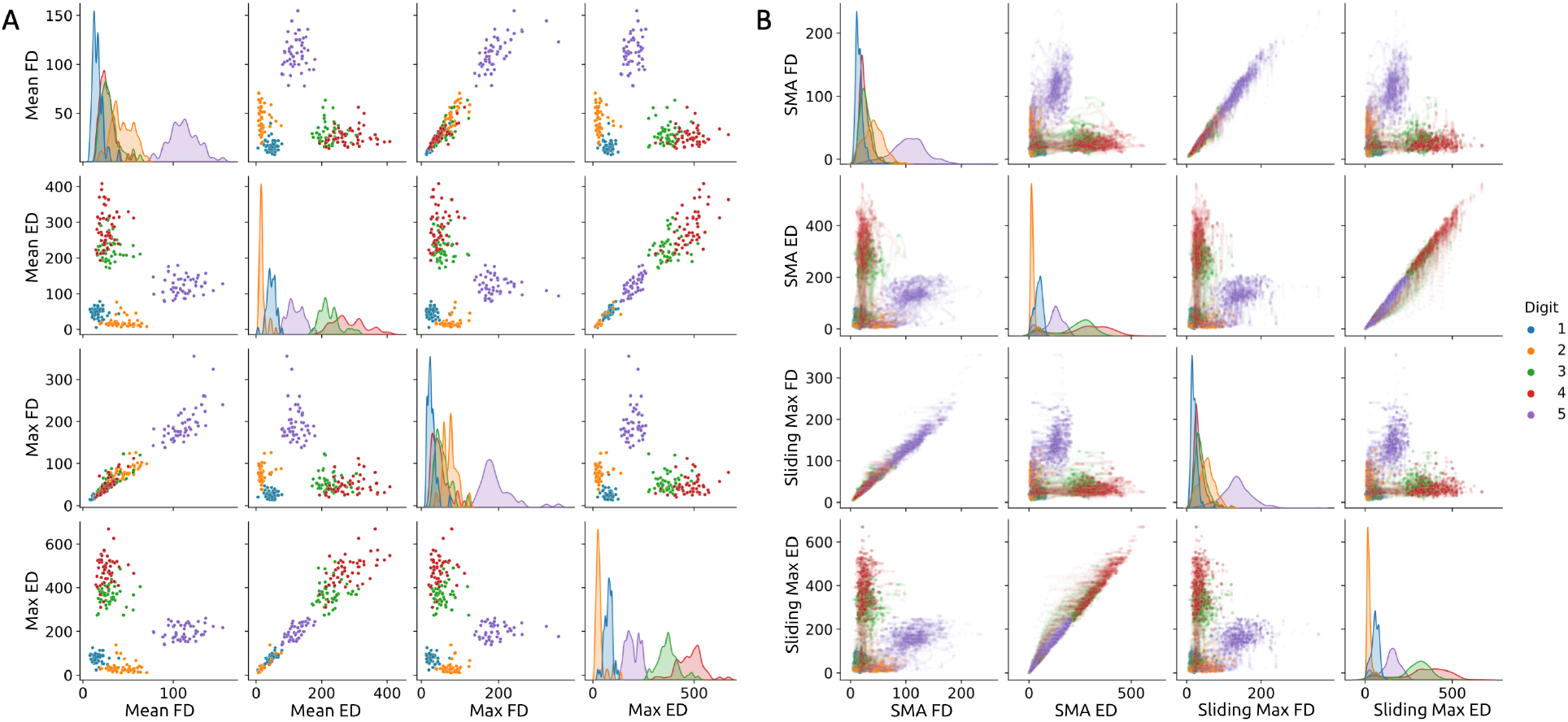
Pairwise plot of all four features against each other (‘mean’ and ‘max’ from both EMG channels) using **(A)** data from the entire trial (1600ms) and **(B)** data from a 280ms long sliding window, during the trial. SMA; smoothed moving average. In panel B, due to the increased number of data points, the transparency level was adjusted to avoid complete occlusion of overlapping points. The diagonal cells show the probability distribution of the respective feature for each digit.

Using a kNN classifier with *k* = 3, a decoding accuracy of 94% was achieved using the total window length, and a 90.4% accuracy using the sliding window length 280ms. A confusion matrix using this sliding window was computed to investigate the source of classification errors (see **Fig 3**), which showed substantial confusion decoding middle and ring finger movements (digits 3 and 4).

**Fig 3:**
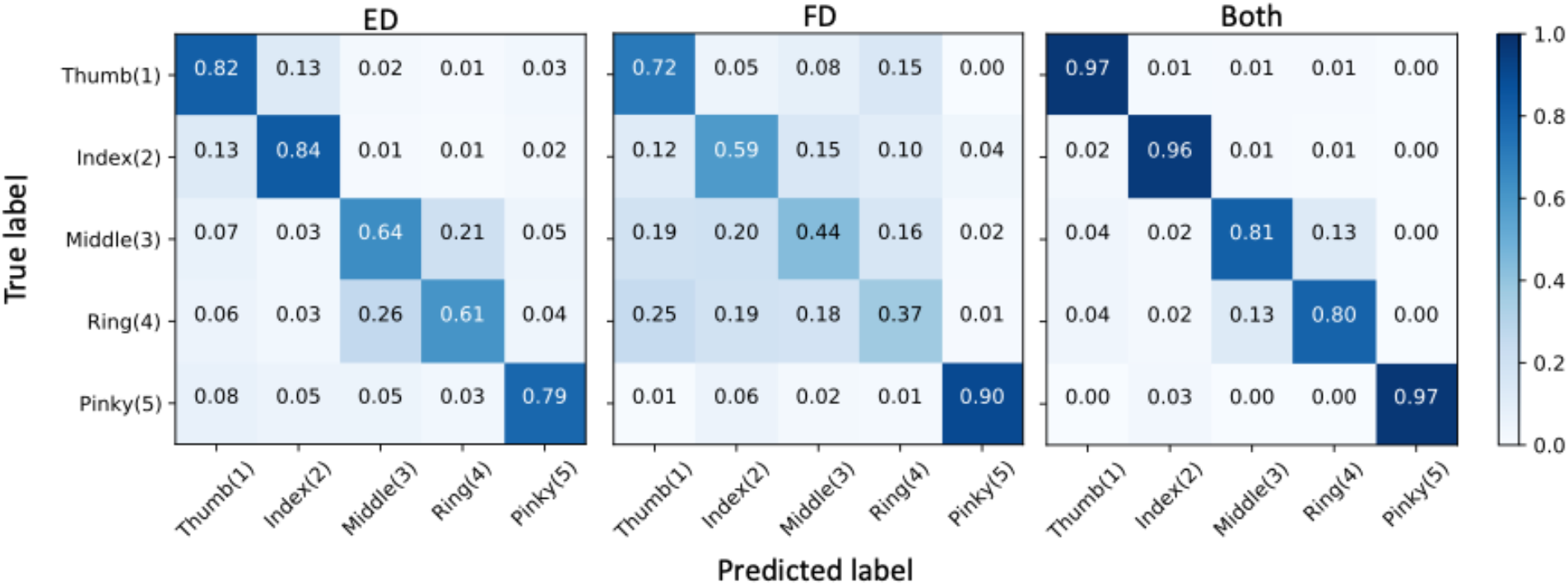
Normalized confusion matrix for each muscle independently, as well as together (ED, FD, Both). Features used were the ‘max’ and ‘mean’ taken from a 280ms sliding window, and tested throughout the 1600ms of the trial using kNN with *k* = 3.

### Unsupervised Learning Approach

We also performed an exploratory investigation into the separability of the five digit classes with an *unsupervised* learning approach. The DBSCAN classifier with standard settings (*epsilon* = 0.5, *minimum samples* = 10) and *k-Means* (with the number of clusters pre-defined as 5) were compared; with *k-Means* being superior when using the ‘max’ feature only (79% of labels correct), and when using the combination of ‘max’ and ‘mean’ (85% of labels correct), **Fig 4.**

**Fig 4:**
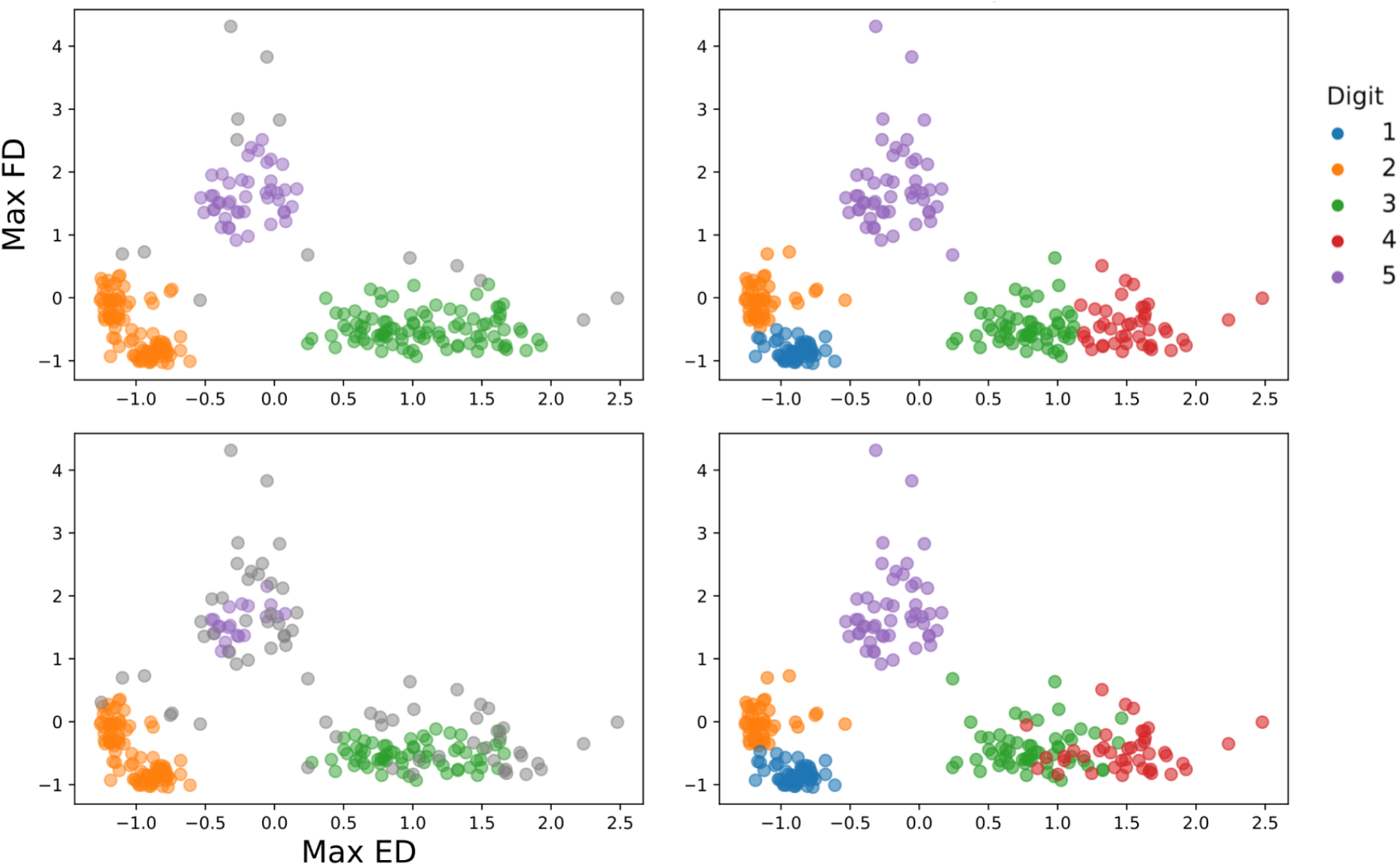
Comparison of unsupervised cluster based machine learning algorithms, DBSCAN (left) and k-Means with 5 clusters (right). (**Top**) classifications using only the ‘max’ of each channel as feature input. (**Bottom**) classifications using ‘max’ and ‘mean’ as feature inputs (only ‘max’ plotted). The x and y-axis scales are arbitrary values. Grey points in DBSCAN represent uncategorized data points (i.e., outliers that are too far from a central node).

## Discussion

The aim of this simple proof-of-concept study was to investigate the feasibility of an EMG decoding approach for use in mixed reality real-time applications. We demonstrate that five different movements corresponding to finger extensions of each individual digit can be decoded with an accuracy >90.4% using only two EMG channels, 280ms sliding windows, and a standard machine learning pipeline that uses simple ‘mean’ and ‘max’ features with a k-nearest neighbors classifier. The entire pipeline is implemented on low-cost off-the-shelf hardware and open-source software. We illustrate how this approach can be used to incorporate an EMG-based interface for human interaction with a virtual environment, such as controlling a user interface (see **Supplementary Material 1**), interacting with a ‘smart’ home (see **Supplementary Material 2**), or in gamified rehabilitation.

One limitation of this study is that we simply chose extension movements of the individual digits, whereas the repertoire of movements could be expanded to include various hand and finger signs and gestures. Also, the set of movements was not optimized for decoding, for example, the confusion between the middle and ring fingers is likely due to the physiological overlap and resultant “enslaving” effects of innervation, as documented by Zatsiorsky et al. (2000). This “enslaving” effect can also be visualized when closer examining the results of the unsupervised machine learning algorithms, as they too failed to adequately separate the two overlapping groups of data from the third and fourth digits. When excluding these two digits, our real-time method reaches closer to 97% accuracy.

Additionally, though we purposefully chose to use only two EMG sensors, we did not optimize the placement of the sensors for the task other than choosing the main muscles for finger extensions (the ED), and contractions (the FD). To this point, the sensor on the ED produces across-the-board higher classification accuracies than the sensor on the FD. This very well could be due to the experimental design, as dorsal extension of the fingers does call for full activation of the ED, yet the subsequent return to a ‘flat’-handed home position would only partially activate the FD (full activation would require more-so closing the digits into a fist). Likely, attuned placement of the sensors in line with specific task requirements, a higher number of EMG sensors, or using a sensor fusion approach by including additional sensors such as computer vision and/or accelerometers would enable an even higher classification accuracy. It also might be necessary when considering applications spanning a greater range of movement classes, especially when they are largely overlapping. Furthermore, using only a single subject is an obvious limitation, but given the nature of similarity in innervation between people (Yu et al. 2004), and that this was a proof-of-principle demonstration, is sufficient for this purpose.

The main conclusion of this study in light of the above limitations, a satisfactory decoding accuracy is feasible in a real-time setting. Moving toward biosignal-controlled devices transitions from the conceptual framework of the use of computers as a tool toward one that is an intuitive extension of oneself and further expands the capabilities of the human experience. We look forward to the next generation of human-computer interfaces that utilize biosignals to enable intuitive and frictionless embodiment for professional, recreational, and medical applications.

## Supplementary Material

Supplementary Material 1: video “EJM_UI”, showing EMG / eye tracking to trigger UI menu

Supplementary Material 2: video “EJM_SmartHome”, showing EMG / eye tracking / gesture recognition to interact with “smart home”

